# Density-dependent expression of epitranscriptomic, stress, and appetite regulating genes in Atlantic salmon

**DOI:** 10.1101/2025.01.19.633827

**Authors:** Morgane Frapin, Laura Quispe, Joana Troka, Jenni M. Prokkola, Ossi Laurikainen, Pekka Hyvärinen, Craig R. Primmer, Tutku Aykanat, Ehsan Pashay Ahi

**Author notes:** Corresponding author: Morgane Frapin. co-senior authors (Tutku Aykanat, Ehsan Pashay Ahi).

## Abstract

Intraspecific competition due to e.g., density, has substantial influence on fitness dynamics and life histories, but the underlying physiological mechanisms are often complex, and the molecular basis unclear. Further, designing laboratory experiments to measure physiological responses that reflect natural conditions is challenging. Here, we reared Atlantic salmon juveniles in semi-wild conditions in two naturally occurring densities to investigate the molecular mechanism of density-related changes in the hypothalamus, a key brain region regulating stress and energy homeostasis. We measured density-dependent changes in the expression of 12 genes involved in appetite and stress regulation, and 16 genes involved in post-transcriptional regulation of gene expression via m^6^A RNA methylation. This gene set includes paralog pairs, to account for potential functional divergence associated with salmonid genome duplication. We also quantified genotype-environment interactions between density and two major life-history loci, *vgll3* and *six6*. We found significant density-related differences in the expression of genes coding for corticotropin-releasing factors, appetite stimulators and inhibitors, and m^6^A RNA methylation actors. Moreover, a paralogue of an appetite inhibitor showed a density-dependent pattern that was the opposite of what was expected. *Six6* locus was also associated with changes in the expression of epitranscriptomic markers, including two writers and one eraser. Our results highlight that individuals’ response to density in natural conditions is shaped by a complex interplay between stress, appetite and epitranscriptomic pathways in the hypothalamus. In addition, the functional divergence of paralogs indicates a potential role of genome duplication shaping such a response. We emphasize the value of integrating different physiological responses at the molecular level to better understand ecological processes affected by environmental change.

## 1. Introduction

Population density is an important factor in ecology, affecting resource availability and social interactions among conspecifics (Nislow et al., 2010; Reznick, 2016). An increase in population density intensifies the intraspecific competition for resources, leading to alterations in energy homeostasis and stress levels, and subsequently impaired growth and reproductive success (Auer et al., 2012; Grossman & Simon, 2020; Reznick, 2016). Fishes are good models for studying density-dependent physiological responses, as density has a significant effect on their growth and survival (Grossman & Simon, 2020; Matte et al., 2020). In the wild, riverine habitats are important ecosystems that are particularly affected by density-dependent processes, due to the limited area and resources, providing a valuable case for studying soft selection and the effects of density changes (Jonsson et al., 1998). For example, the strong territorial behavior of Atlantic salmon to occupy favorable habitats can be beneficial in poor habitats, but may not be energetically efficient in resource rich habitats (Blanchet et al., 2006; Jonsson et al., 1998; Nislow et al., 2010). In laboratory or hatchery settings, fish are reared at much higher densities, and competition and food acquisition dynamics do not reflect the natural conditions. This is likely limiting individuals from displaying natural behaviors. As such, two habitats (*wild versus* laboratory conditions) are likely to result in different physiological responses to density changes (Blanchet et al., 2006; Höjesjö et al., 2004).

An individual’s response to density may invoke multiple physiological and/or stress pathways, which can result in complex responses at the population level (Birk et al., 2020; Hodgson et al., 2017), with potential impacts on life histories (Archer et al., 2020). At the organismal level, such a response is regulated by hypothalamus, which exhibits distinct molecular pathways for various environmentally induced responses (Schwartz et al., 2000; Smith & Vale, 2006). The hypothalamus is a key region of the brain that connects the central nervous system to the endocrine system. It regulates feeding behavior by balancing appetite-stimulating genes (orexigenic genes), such as Agouti-related protein (AgRP) and neuropeptide Y (NPY), and appetite-suppressing genes (anorexigenic genes), such as cocaine- and amphetamine-regulated transcript (CART) and pro-opiomelanocortin (POMC). Together, these mechanisms help the body regulate energy intake and expenditure in response to nutritional status (Parker & Bloom, 2012). The hypothalamus is also functioning as the primary interface for stress detection by orchestrating the glucocorticoid receptor signaling pathway and initiating the secretion of corticotropin-releasing hormone/factor (CRH or CRF). This activates the hypothalamic-pituitary-adrenal axis, leading to the release of cortisol, a key stress hormone that prepares the body to respond to stressors (Smith and Vale 2006). Additionally, the hypothalamus plays a pivotal role in sexual maturation by releasing gonadotropin-releasing hormone, influencing the onset of puberty and reproductive capacity (Sisk & Foster, 2004). These regulatory roles of the hypothalamus are conserved across vertebrates, underscoring its fundamental importance in maintaining homeostasis and enabling responses to environmental changes and internal conditions (Denver, 2009). Thus, understanding hypothalamic regulation may be critical in partitioning the relative contribution of different physiological processes, such as stress and appetite pathways, which may help to pinpoint ecological functions that underlie density-dependent dynamics in the wild.

Emerging research highlights the role of m^6^A RNA methylation in modulating hypothalamic signals related to feeding, stress, and sexual maturation. Thus, m^6^A RNA methylation may play a role in density-dependent regulation by controlling gene expression in response to environmental changes. m^6^A RNA methylation is a widespread, reversible, and evolutionarily conserved mRNA modification (Ahi & Singh, 2024; Shi et al., 2022). The addition of m^6^A RNA methylation is facilitated by the m^6^A methyltransferase complex, which involves writer proteins (such as Mettl3, Mettl14, and Wtap), and removal is carried out by eraser demethylase proteins (such as Fto and Alkbh5) (Zaccara et al., 2019). Reader proteins (e.g., Ythdf1/2/3 and Ythdc1/2) recognize the m^6^A modifications, directing the methylated mRNA towards various outcomes, including degradation, stabilization, transportation, and either promotion or inhibition of translation (Liao et al., 2018; Zhao et al., 2017). Investigating whether m^6^A RNA modification affects hypothalamic signaling that controls stress and feeding in response to environmental changes, such as population density, could provide insight into growth flexibility and the proximal mechanism of phenotypic plasticity in changing environmental conditions (Jonsson et al., 1998).

The Atlantic salmon (S*almo salar*) is a culturally and economically important anadromous fish species that spends its juvenile phase in the freshwater. The freshwater phase is highly influenced by density effects (Thorstad et al., 2010). While molecular and/or physiological response to density have been widely investigated in Atlantic salmon reared in artificial conditions (Liu et al., 2015; Sundh et al., 2019; Sveen et al., 2016; Y. Wang et al., 2019), these hardly reflect natural settings, and conclusions from these studies can’t necessarily be extrapolated to wild conditions. Studying the molecular/physiological pathways of density regulation is feasible in Atlantic salmon in semi-natural environment, and therefore more natural-like physiological responses can be obtained. Furthermore, Atlantic salmon may also be a good model to study molecular mechanisms of life history variation and potential interplay with hypothalamic regulation of density. Genetic variants located by *vgll3* gene in chromosome 25 and *six6* gene in chromosome 9 have been associated with a sea age at maturity in salmon (Barson et al., 2015; Sinclair-Waters et al., 2020). This simple genetic architecture is also associated with various behavioral and physiological pathways (e.g., Aykanat et al., 2020; Bangura et al., 2022; Prokkola et al., 2022), and a recent study reported that the hypothalamic expression of genes involved in m^6^A RNA methylation is influenced by *six6* locus, sex and maturation status in Atlantic salmon (Ahi et al., 2024). As such these genes may partly be associated with hypothalamic endocrine pathways that respond to environmental, dietary, and physiological cues (Gerisch & Antebi, 2011; Stearns, 2011).

In this study, our primary aim is to investigate how changes in density affect fundamental molecular physiological processes in an environment that closely mimics natural conditions. We explore the hypothalamic expression of genes related to appetite regulation, stress hormone signaling pathways and m^6^A RNA methylation in 2-year old Atlantic salmon parr reared in semi-natural stream conditions, and naturally fed on mainly benthic prey items, at two different densities similar to those found naturally in the wild (Finstad et al., 2009). In addition, the presence of the genetic variation at the two loci associated with sea age at maturity, *vgll3* and *six6*, allowed us to assess their effect, as well as their interaction with density, on the gene expression of the targeted hypothalamic pathways. Given the whole-genome duplication which occurred approximately 80 million years ago in salmonids (Dysin et al., 2022; Lien et al., 2016), we also investigate potential functional evolution of the duplicated genes by characterizing the expression pattern of paralogs (Force et al., 1999; Kassahn et al., 2009).

## 2. Materials and Methods

### 2.1. Fish rearing

All fish experiments were conducted in accordance with the Finnish Project Authorisation Board approval (ESAVI/4511/2020). In this experiment, Atlantic salmon were obtained from broodstocks reared by the Natural Resources Institute Finland (LUKE, Laukaa, Finland). The parents of the broodstocks, i.e., the grandparents of the experimental fish originated from the Kymijoki River in Southern Finland. From the broodstock, three unrelated males and females heterozygous for the alleles associated with earlier (E) and later (L) maturation at both the *vgll3* and *six6* loci were crossed in October 2019 to obtain three unrelated families each containing offspring with all combination of genotypes for these loci. Fertilized eggs were incubated in the dark at 7°C at the University of Helsinki until the alevin stage as described earlier (Debes et al., 2021), and then transferred late January 2020 to the Kainuu Fisheries Research Station of LUKE in Paltamo (Finland). The fish were reared in a seasonal temperature and daylight cycle and fed based on their size (Veronesi vita 0.2-0.8mm, Raisioagro Oy) in a flow-through system supplied with water from a nearby lake. In January 2021, fish were PIT-tagged (12 mm) and small fin clips were taken for genotyping individuals for sex and *vgll3* and *six6* loci (Sinclair-Waters et al., 2022). In May 2021, a subset of fish was released in six parallel outdoor channels (length: 25 meters, width: 1.5 meters, approx. 39 m^2^) with concrete walls and bottoms covered by 15 cm layer of gravel-to-pebble-sized particles (8-50 mm in diameter). These semi-natural stream channels were fed by water from the same lake as the indoor tanks to maintain a flow rate of 0.15-0.25 ms^−1^. The fish were naturally fed on mainly benthic prey items from the bottom substrate (insects, arachnids, and crustaceans), but also some surface prey (such as adult midges, mosquitoes), similar to their natural habitat. All fish selected in this experiment were smolts based on silvery coloration. Each of the three families was divided between two stream channels with two density treatments, which initially had 48 (1.2 fish/m^2^) and 112 (2.8 fish/m^2^) fish per low-density and high-density treatment streams, respectively. However, an infection outbreak caused by *Saprolegnia spp.* right after the transfer to the outdoor tanks inflicted high mortalities to the experimental fish (as high as 46 % mortality in less than a month). To bolster the density effect, high-density tanks were supplemented with additional fish, which were not included in subsequent analysis. These additional fish originated from the same families and were reared in the same indoor tanks as the experimental fish prior to the transfer to the natural stream channels. In August 2021, due to practical reasons, all fish were transferred from the channels to six circular stream channels (Outer circumference: 26.15 meters, width: 1.5 meters, approx. 39.2 m^2^) with concrete walls and a bottom covered with a 15 cm layer of gravel (8-50 mm in diameter) and a gravity driven flow of 0.11 ms^−1^. The densities by the end of the experiments ranged from 0.35-0.51 fish/m^2^ and 1.62-2.05 fish/m^2^ for low-density and high-density stream channels, respectively (see Supplementary table 1 for details).

### 2.2. Sampling and RNA extraction

On 25-27 April 2022, fish were caught from the streams by netting after water level in each stream was lowered and placed in aerated 40-L buckets until sampling the same day. Fish were euthanized with an overdose of benzocaine (200 mg/L), weighed, and measured. The hypothalamus of fish from both densities and all genotypes were sampled, snap-frozen in liquid nitrogen and stored at −70°C prior to processing.

To avoid introducing variation associated with maturation status, only immature individuals were selected for this study. Maturity was assessed by checking the presence of milt in males (females are immature at that age). The hypothalamus of 81 individuals (40 males and 41 females) was selected. The individuals were either homozygous (EE or LL) or heterozygous (EL) for *six6* and *vgll3* loci (N = 3 - 10 samples per genotype, sex and density) (see, Supplementary table 2 for details). Total RNA was extracted using Nucleospin RNA kit (Machery-Nagel). Homogenization and cell lysis steps were performed simultaneously by transferring the frozen hypothalamus in a 2 mL screw cap tube containing 1.4 mm ceramic beads (OMNI), 3.5 μL of DTT 1M and 350 μL of RA1 buffer and placing the tubes in a Bead Ruptor Elite (OMNI) at 4 ms^−1^ for 3×20 seconds with a 10 second break in between. The subsequent steps followed the manufacturer’s instructions and RNA was eluted in 40 μL of RNase-free water. RNA quantity and quality were assessed using Nanodrop 2000 (Thermo Fisher Scientific) and agarose gel electrophoresis.

### 2.3. cDNA synthesis and qPCR

Synthesis of cDNA was performed using iScript cDNA Synthesis Kit (BioRad) with 1 μg of RNA. The genes *ef1a* and *hprt1* were used as housekeeping genes (Ahi et al., 2024). The expression of six genes related to stress (*nr3c1*, *nr3c2*, *crf1a1*, *crf1a2*, *crf1b1*, *crf1b2*), six genes related to appetite regulation (*agrp1*, *cart2a*, *cart2b*, *pomca1*, *pomca2*, *npya1*) and 16 genes related to epitranscriptomic mechanisms (*mettl3*, *mettl14*, *wtap*, *alkbh5-1*, *alkbh5-2*, *fto-1*, *fto-2*, *ythdc1-1*, *ythdc1-2*, *ythdc2*, *ythdf1-1*, *ythdf1-2*, *ythdf1-3*, *ythdf2-1*, *ythdf2-2*, *ythdf3*) were measured using quantitative real-time PCR (qPCR) (genes and primer sequences are described in Supplementary table 3). The qPCR reactions were performed in a final volume of 10 μL containing 1 μL of cDNA diluted 1:10 in nuclease-free water, 250 nM of forward and reverse primers and 5 μL of PowerUp™ SYBR™ Green Master Mix (ThermoFisher Scientific). The qPCRs were performed on 384-well plates using the BioRad CFX384 Real Time PCR System, with each plate measuring the expression of a single target gene across all samples that were present in triplicate. The Cqs (quantification cycle) were obtained with a manually fixed threshold located in the exponential part of the amplification. The qPCR program and primer efficiency calculation were performed as described in Ahi, Richter, et al. (2019). For each sample, the Cq values of the triplicates obtained for each gene were averaged. The ΔCq values were calculated by subtracting the geometric mean of the Cq values of the housekeeping genes, *ef1a* and *hprt1*, from the Cq values of the target genes. Then, for each gene, the ΔCq values were subtracted by the ΔCq value of the sample with the lowest expression to obtain the ΔΔCq. The relative quantifications (RQ) were calculated as 2^−AACq^ and statistical analyses used the log_2_ RQ (Livak & Schmittgen, 2001).

### 2.4. m^6^A RNA methylation quantification

Quantification of m^6^A RNA methylation was performed on the 81 total RNA extracted from the hypothalamus using the EpiQuikTM m6A RNA Methylation Quantification Kit (Epigentek). Since one preparation kit can accommodate a maximum of 96 tests, two kits were used to include all samples in duplicate, along with negative and positive controls. Samples were randomly assigned between the two kits to account for potential batch effects in the statistical model. RNA samples were diluted to 33.33 ng/μL in RNase-free water, and the concentration was verified with Nanodrop 2000 (Thermo Scientifc). m^6^A RNA methylation quantification was performed according to the manufacturer’s protocol with 6 μL of the diluted total RNA (approximately 200 ng). Negative and positive controls from the kit were included in each plate. The absorbance was then read at 450 nm with the EnSpire Multimode Plate Reader (PerkinElmer). m^6^A percentages were calculated according to the manufacturer’s protocol based on the standard curve and with the amount of total RNA input adjusted based on the actual concentrations of the dilutions used.

### 2.5. Statistical analysis

All the statistical analysis were done in R (version 4.2.2). A linear mixed effect model was fitted using lmer function in the *lme4* package (version 1.1-31) (Bates et al., 2015), with REML=FALSE option, using a general formula as below:

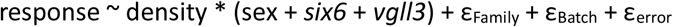

In which, response is either *body mass, gene expression or m^6^A %.* Density was modelled as a fixed effect interacting with sex, *six6* and *vgll3* genotypes. Genotypes were coded categorically to account for non-additivity. e_Family_, e_Batch_ are error terms associated with family and batch effect, and e_error_ is the residual normal variation. e_Batch_ was only included for the *m^6^A % response* to account for the kit effect. The significance of the fixed effects terms was estimated using type 1 F-test using the anova function in the stats package (version 4.2.2) in R. The normal distribution of the model residuals was checked using the simulateResiduals function in the *DHARMa* package (version 0.4.6, (Hartig et al., 2024). P-values for the gene expression analysis were adjusted for multiple comparisons per model term (i.e., density, sex, *six6* genotype, *vgll3* genotype, and the interaction terms: density x sex, density x *six6*, density x *vgll3*) using the Benjamini-Hochberg method to control the false discovery rate (Benjamini & Hochberg, 1995). An adjusted p-value below 0.05 was considered statistically significant. For terms with significant effects, post-hoc pairwise comparisons were performed using estimated marginal means (EMMs) via the *emmeans* package (version 1.8.2, (Lenth et al., 2024). These pairwise comparisons allowed us to examine specific contrasts between the levels of significant variables. The p-values presented in the graphs are the adjusted p-values, obtained from these post-hoc pairwise comparisons and corrected using the Benjamini-Hochberg method. The p-values in the text correspond to the adjusted p-values obtained after adjusting the ANOVA results for the false discovery rate.

To investigate potential interactions between the target genes, body mass, and m^6^A percentage, we performed a correlation analysis. Pearson correlations were calculated using cor.test function (*stats* package version 4.2.2) on the residuals of the fitted body mass, gene expression and m^6^A percentage models. This analysis was carried out separately for the high- and low-density groups.

## 3. Results

### 3.1. Fish body mass was influenced by density treatment

After one year in different densities, fish body mass was significantly lower in high-density (p<0.001; F_1;78.37_=80.425) (Figure 1). However, neither sex nor the *six6* and *vgll3* genotypes had a significant effect on fish body mass.

**Figure 1:**
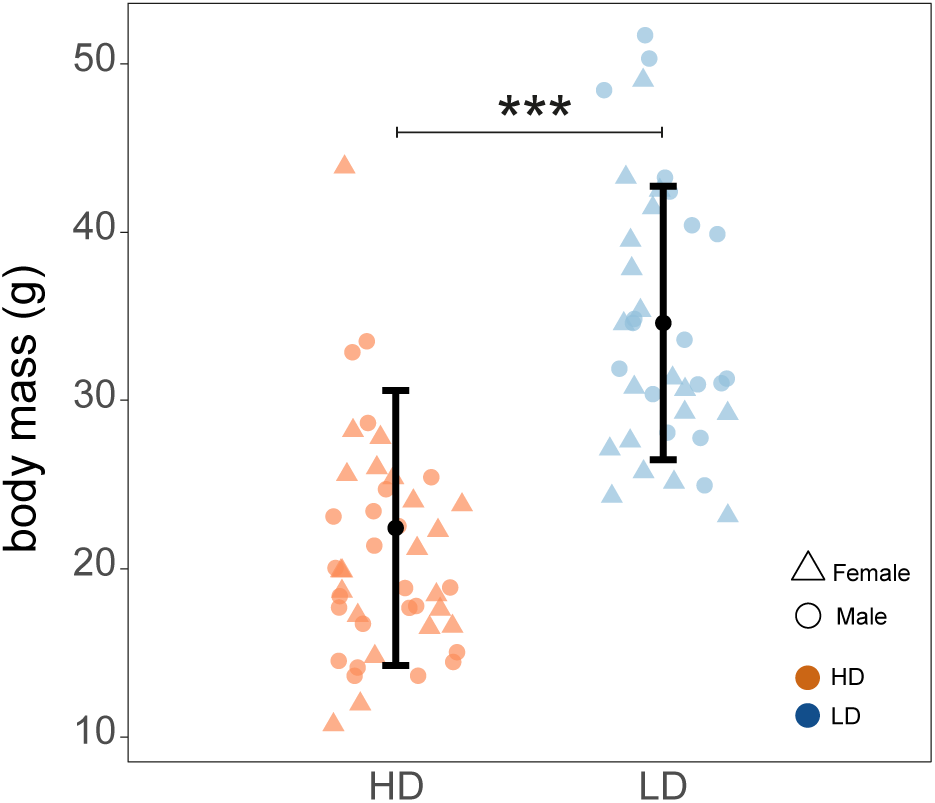
Fish body mass (in grams) measured after one year under density treatment. Each dot represents the body mass of an individual fish in the high-density (HD) group (n = 44, orange dots) and low-density (LD) group (n = 37, blue dots). The black points display the estimated marginal means for each treatment group. Error bars indicate 95% confidence intervals. ***: p-value<0.001.

### 3.2. Density and genetic factors affect expression levels of Corticotropin-Releasing factor genes

Density, and *six6* and *vgll3* genotypes influenced the expression of corticotropin-releasing factor (*crf)* genes but not the expression of glucocorticoid receptor (*nr3c*) genes (Supplementary table 4). The level of expression of *crf1a1* was significantly higher in the high-density treatment (p=0.025; F_1;79.26_=8.598) while *crf1b1* expression was significantly lower (p=0.003; F_1;81_=13.801). Moreover, *crf1a1* expression was also significantly influenced by *six6* genotype (p=0.018, F_2,78_=6.78) with a higher level of expression in *six*6*EE (95% CI=1.64-2.66) compared to *six6*\*EL genotype individuals (95% CI=0.84-1.91). An interaction effect between density and *vgll3* genotype was found for one of the corticotropin-releasing factor paralogs, *crf1b2* (p=0.039, F_2,79_=7.15), with a higher level of expression of the gene in *vgll3*\*LL in low-density (95% CI=1.02-1.43) compared to *vgll3*\*EE (95% CI=0.44-0.91) and *vgll3*\*EL (95% CI=0.49-1.02), and compared to *vgll3*\*LL in high density (95% CI=0.51-0.91, Figure 2).

**Figure 2:**
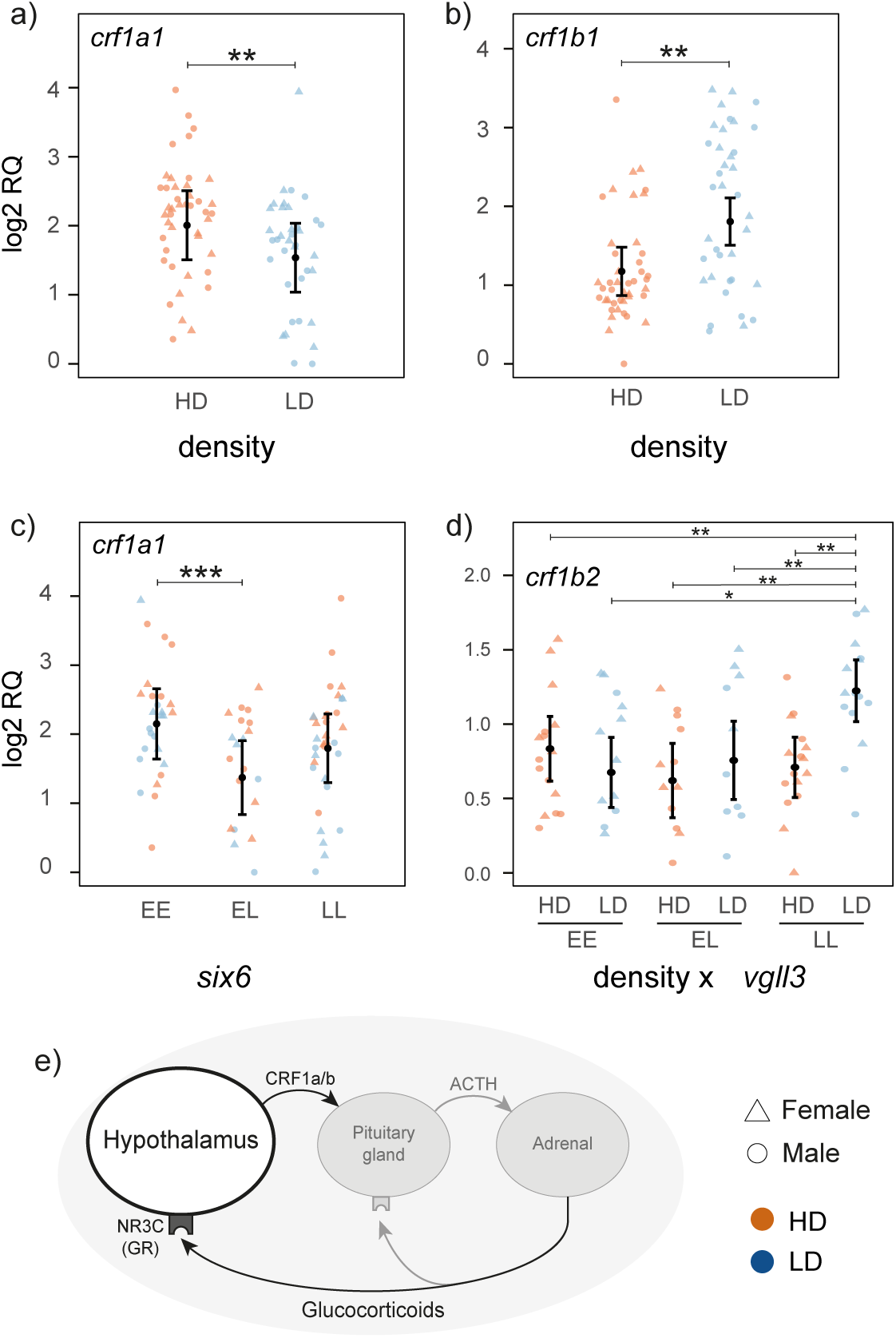
Relative mRNA expression (log2RQ) level of stress-related genes affected by density **(a,b)**, six6 **(c)** and the interaction between vgll3 and density **(d)**. Error bars indicate 95% confidence intervals. *: p-value<0.05; **: p-value<0.01; ***: p-value<0.001. **(e)** Diagram summarizing the implication of the studied genes in stress regulation. Stress is sensed by the hypothalamus. This triggers a hormonal cascade initiated by the release of corticotropin-releasing hormone/factor (CRH/CRF). Secretion of CRH stimulates the release of adrenocorticotropic hormone (ACTH) into the bloodstream, which then acts on the interrenal cells located in the fish head kidney, stimulating the release of cortisol (Gorissen & Flik, 2016). Cortisol binds to glucocorticoid or mineralocorticoid receptors present in various tissues, e.g. NR3C, leading to physiological and metabolic adaptations in response to the stressor and affecting fish growth (Barton & Iwama, 1991; Takahashi & Sakamoto, 2013)

### 3.3. Density influences the expression of both anorexigenic and orexigenic pepting-coning genes

The expression levels of genes encoding both appetite-stimulating (Agrp) and appetite-inhibiting (Cart2a/b) peptides were affected by density treatment (Figure 3) (Supplementary table 4). The level of expression of *agrp1* was significantly lower at low-density (p<0.001; F_1;79_=51.64). The paralogs *cart2a* (p=0.05; F_1;81_=6.51) and *cart2b* (p=0.004; F_1;81_=12.67) showed opposite patterns, with *cart2a* showing higher expression and *cart2b* lower expression at low-density. The level of expression of the *cart2a* was also influenced by *six6* genotype (p=0.006; F_2;81_=9.38), where heterozygote (95% CI=0.81-1.43) exhibited significantly lower expression compared to homozygotes (95% CI=1.62-2.15 and 1.31­*1.77* for *six6*\*EE and *six6*\*LL, respectively).

**Figure 3:**
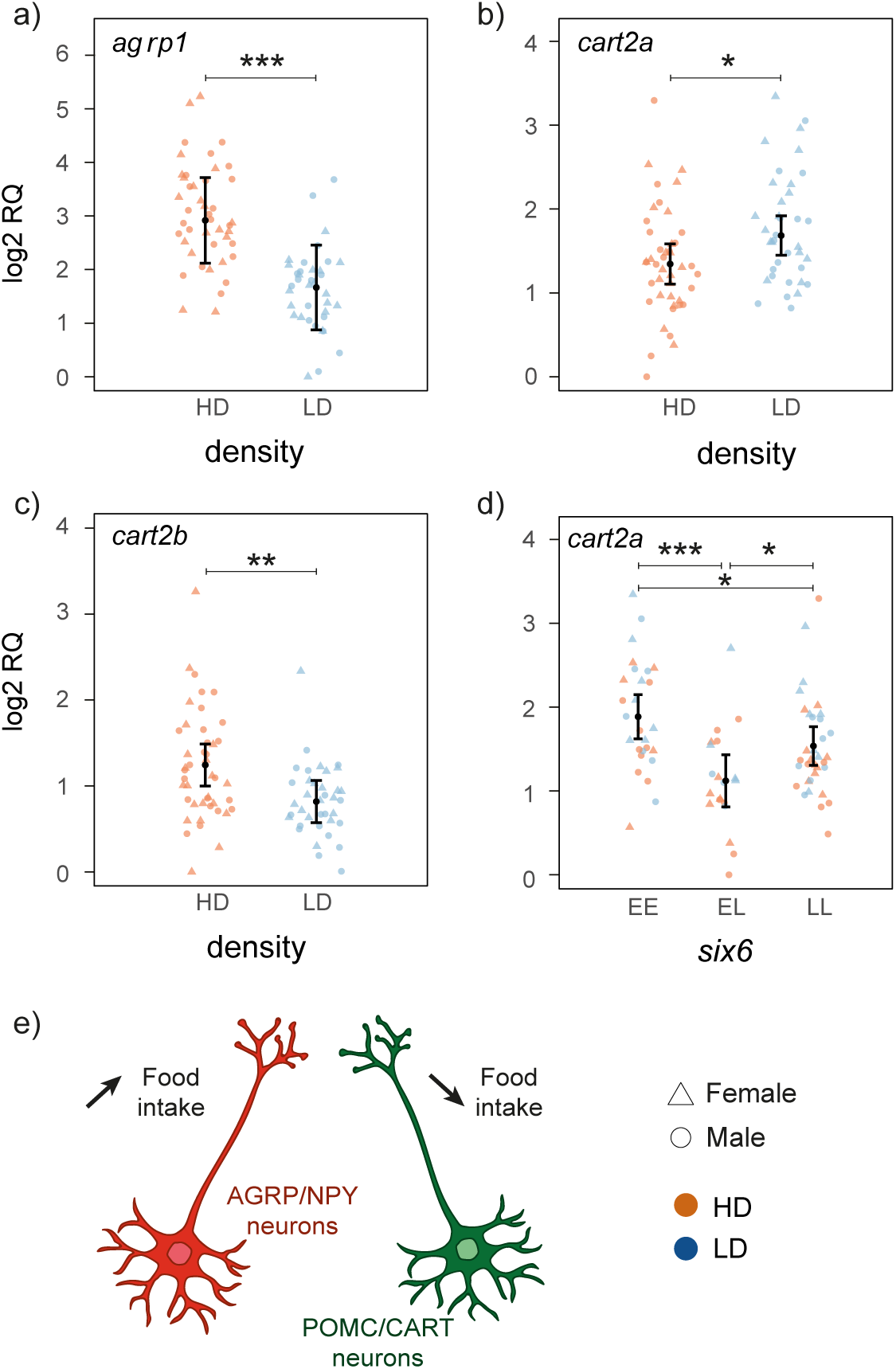
Relative mRNA expression (log_2_RQ) level of appetite-related genes affected by density **(a,b,c)** six6 **(d)**. Error bars indicate 95% confidence intervals. *: p-value<0.05; **: p-value<0.01; ***: p-value<0.001. (e) Diagram summarizing the implication of the studied genes in the appetite regulation. Orexigenic (AGRP and NPY) and anorexigenic (POMC and CART) neurons in the arcuate nucleus of the hypothalamus detect the presence of hormones (insulin, leptin) in the bloodstream and induce an increase (orexigenic response) or a decrease (anorexigenic response) in food intake respectively (Schneeberger et al., 2014).

### 3.4. Density and *six6* genotype affect the expression of genes encoding m^6^A regulators without altering overall m^6^A levels

Both density and *six6* genotype influenced the expression of genes encoding m6A RNA methylation writers (Supplementary table 4). The level of expression of *mettl3* (p=0.034; F_i;8i_=7.55) and *ythdf1.2* (p=0.003; F_1;80_=14.82) was high in low-density group. The *six6* genotype influenced the expression levels of two writers, *mettl3* (p=0.006; F_2;8i_=8.52) and *wtap* (p=0.031; F_2;81_=5.81). For *mettl3,* expression was lower in *six6*\*EL (95% CI=0.35-0.62) compared to *six6*\*LL (95% CI=0.69-0.89). *wtap* expression was reduced in *six6*\*EL (95% CI=0.66-1.04) compared to both *six6*\*EE and *six6*\*LL (95% CI=1.05-1.38 and 1.01-1.30 for *six6*\*EE and *six6*\*LL, respectively). The level of expression of /tol, an m^6^A eraser, was also significantly affected by the *six6* genotype (p=0.041; F_2;78_=5.25) with a lower level of expression in *six6*\*EL (95% CI=0.58-1.25) compared to either of the homozygous genotypes (95% CI=0.84-1.52 and 0.81-1.48 for *six6*\*EE and *s/x6*\*LL, respectively, Figure 4). The analysis of the percentage of m^6^A in total hypothalamic RNA showed no significant effects for density (p=0.834; F_1;80_=0.04), sex (p=0.832; F_1;78_=0.05), *six6* (p=0.054; F_2;79_=3.02) and genotypes.

**Figure 4:**
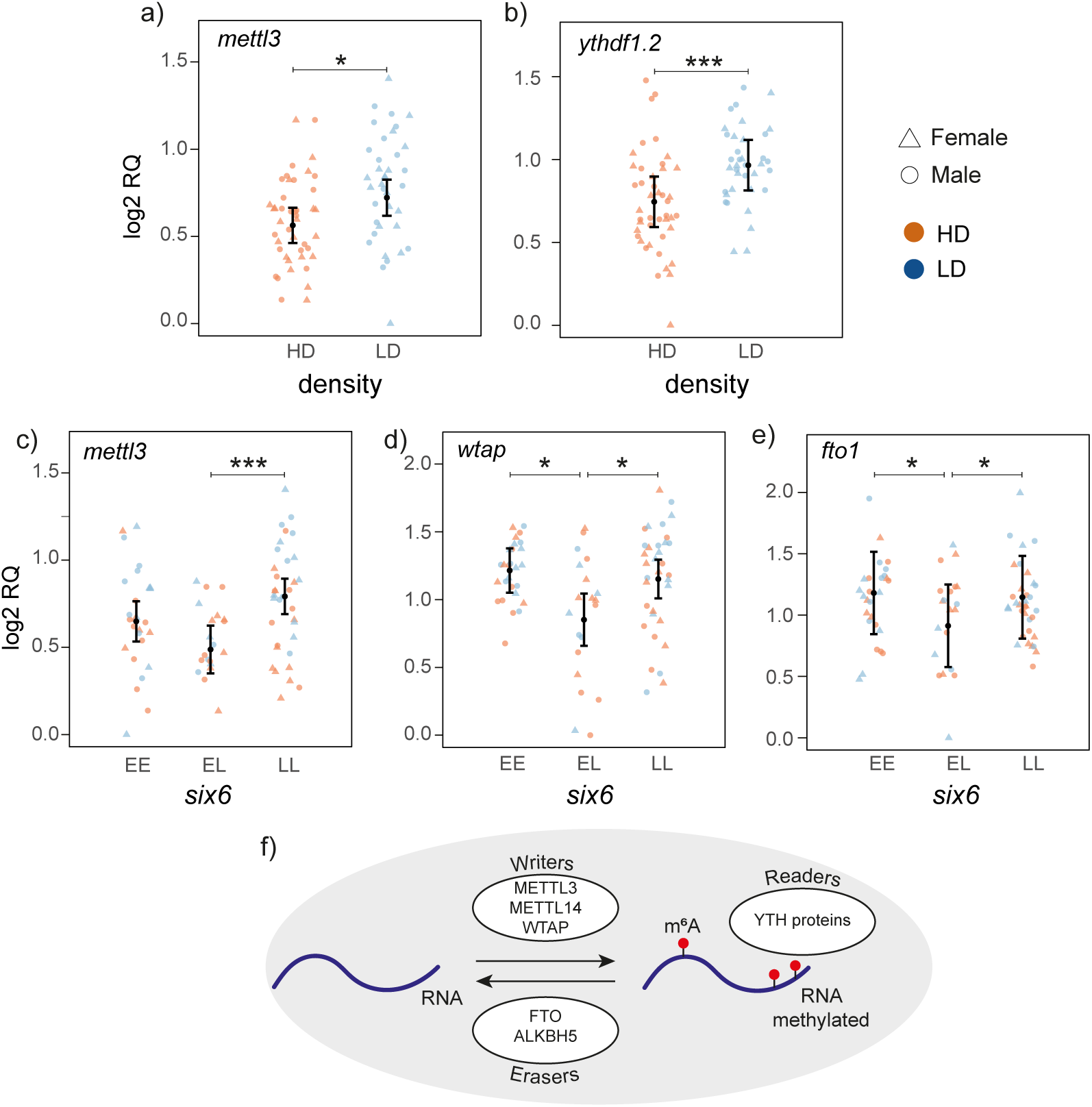
Relative mRNA expression (log_2_RQ) level of m^6^A RNA methylation-related genes affected by density **(a,b)** and six6 (c,d,e). Error bars indicate 95% confidence intervals. *: p-value<0.05; **: p-value<0.01; ***: p-value<0.001. (f) Diagramm summarizing the implication of the studied genes in m^6^A RNA methylation regulation. m6A RNA methylation is added by writers (METTL3, METTL14, WTAP), forming a complex and removed by erasers (FTO and ALKBH5). The fate of RNA is determined by readers (such as the YTH family of proteins) that direct RNA to be translated, transported or degraded (Zaccara et al., 2019).

### 3.5. Correlation between gene expression levels and m6A RNA methylation quantity

The expression levels of *nr3c1* and *nr3c2* were significantly correlated with several genes related to m6A RNA methylation in both high-density (Supplementary figure 1) and low-density (Supplementary figure 2). Furthermore, *nr3c1* was negatively correlated with m6A RNA methylation percentages only in the low-density group (R=-0.48 (p=0.002)) and strongly positively associated with the level of expression of *fto1* in both densities (R=0.7 (p<0.001) in high-density and R=0.8 (p<0.001) in low-density, Figure 5). In the low-density treatment, *cart2b* was negatively correlated with several m6A readers (*ythdc1.2*: R=-0.72 (p<0.001); *ythdc2.1*: R=-0.6 (p<0.001); *ythdf2.2*: R=-0.4 (p=0.01); *ythdf3.1*: R=-0.58 (p<0.001)) but was strongly positively correlated with *alkbh5.1* (R=0.74 (p<0.001)). No strong correlation was found between *alkbh5* paralogs, *alkbh5.1* and *alkbh5.2*, however in low-density condition these two genes showed an opposite correlation with m6A RNA percentage (*alkbh5.1*: R=0.1 (p=0.50) in high-density and R=0.38 (p=0.02) in low-density; *alkbh5.2*: R=0.11 (p=0.48) in high-density and R=-0.38 (p=0.02) in low-density)

**Figure 5:**
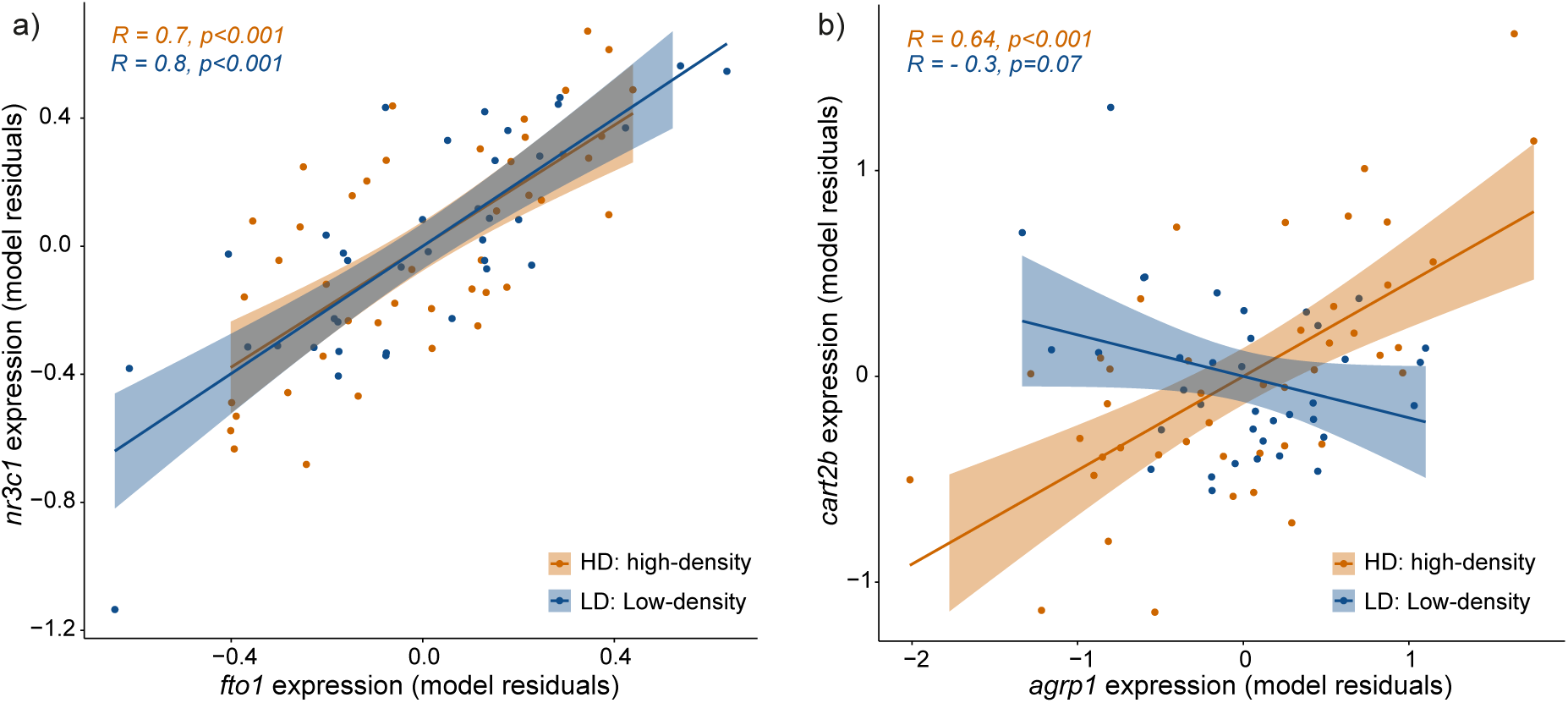
Correlation plots of residual expression levels at different densities **(a)** between fto1 (m^6^A-related gene) and nr3c1 (stress-related gene), and **(b)** between agrp1 and cart2b (appetite-related genes). R: Pearson correlation coefficient.

A positive correlation between two appetite related genes, *agrp1* and *cart2b,* was observed exclusively in the high-density group (R=0.64 (*p*<0.001), Figure 5). These two genes were also negatively correlated with body mass in the high-density treatment (*agrp1*-body mass: R=-0.56 (p<0.001); *cart2b*-body mass: R=-0.62 (p<0.001)).

## 4. Discussion

Individuals can respond to changes in population density by adjusting appetite and stress levels (Nislow et al., 2010; Reznick, 2016), both of which would be regulated via the hypothalamus. The degree of their response to density may also influence growth and, consequently, life-history traits such as timing of maturation. However, little is known about how changes in density affect related molecular pathways in the wild. In this study, we assessed, in a semi-natural environment, the effects of population density on the expression levels of genes related to stress and appetite, and m^6^A RNA methylation in the hypothalamus of Atlantic salmon, in relation to the genotypes of two loci associated with age at maturity (*six6* and *vgll3*). We found that density and *six6* genotype influenced the expression of several genes related to stress, appetite and m^6^A RNA methylation. A *vgll3* genotype effect was also found on a stress-related gene (*crf1b2*) in low-density condition. Our results also suggested widespread expression differences in paralogue gene pairs, which suggests functional divergence in these molecular pathways since the recent genome duplication event (Lien et al., 2016).

### 4.1. Density effects on stress-related genes suggest functional divergence of the *crf* paralogs

Since population density may influence stress levels in fish by affecting food availability and social interactions, we tested the effect of density on the expression of genes encoding corticotropin-releasing factor, CRF, and glucocorticoid receptors, NR3C, which are important ubiquitous stress modulators. We found paralog-specific expression differences; while *crf1a1* was upregulated in the high-density, *crf1a2* expression was not altered, and *crf1b1* was upregulated in the low-density group. Furthermore, in contrast to the two *crf1b* paralogs, the expression of the two *crf1a* paralogs was not significantly correlated. Previous research in aquaculture stocks has shown that the expression of *crf* increases significantly under stress conditions in the hypothalamus of rainbow trout (Conde-Sieira et al., 2010) and in the brain of the Senegalese sole, *Solea senegalensis,* (Wunderink et al., 2011). Interestingly, some studies have shown the opposite trend, with a decrease in *crf* expression under stress conditions, for example in the whole brain of zebrafish exposed to acute restraint stress (Ghisleni et al., 2012) and in a cichlid fish (*Astatotilapia burtoni*) after a month of social stress (Chen & Fernald, 2008). This could be due to a different response depending on the stress conditions, but also to different functions of the *crf* paralogs given that the paralogue studied in teleost fish is not always clearly stated (Grone & Maruska, 2015). Indeed, a recent study showed that all the *crf* paralogs (*crf1a1/2*, *crf1b1/2*) found in Atlantic salmon do not exhibit the same changes in expression patterns under stress conditions, suggesting paralog specificity (Lai et al., 2021). Lai et al., (2021) showed that hypothalamic *crf1a1* expression increased in response to hypoxia and/or hunting stress, in a pattern similar to serum cortisol levels, while *crf1a2* expression remained unchanged. The expression of the *crf1b1* and *crf1b2* paralogs was only altered during the combination of these two stress stimuli (Lai et al., 2021). This potential functional divergence of the paralogs may indicate that certain paralogs are more closely associated with the anorexigenic function previously described in teleosts (Bernier, 2006), while others have a function more related to stress response.

### 4.2. *cart2b* may have an orexigenic effect in high-density conditions

The effect of population density was also investigated on genes involved in appetite regulation, including those encoding orexigenic peptides that stimulate appetite (*agrp* and *npy*), as well as anorexigenic peptides that inhibit appetite (*pomc* and *cart*). We found that *agrp1* showed the strongest response to density, with a highly significant, approximately 2-fold increase in expression at high-density compared to low-density. This might be due to its orexigenic role in fish and/or its responsiveness to stressors. As in many vertebrates, a*grp* has orexigenic effects in several fish species and its expression is induced in response to lack of food availability (Ahi, Brunel, et al., 2019; Cerdá-Reverter & Peter, 2003; Wei et al., 2013). Recent studies of Atlantic salmon have also shown that the hypothalamic expression levels of *agrp1* was increased after exposure to fasting (Kalananthan et al., 2020, 2023), suggesting a similar orexigenic role in salmonids. On the other hand, the expression of *agrp1* also significantly increased in response to stress in zebrafish (Cortés et al., 2018). Therefore, the strong induction of *agrp1* in high density conditions might be an orexigenic response due to lower food availability as well as its response to the stress related to inter-individual competition in higher densities.

We found paralog-specific responses of *cart* to density; *cart2b* had a significantly higher expression in high-density, whereas *cart2a* showed an increased expression in low-density. In zebrafish and many other teleost fishes, *cart* plays anorexigenic role (Volkoff, 2016), however, a study in zebrafish suggested that not all *cart* genes follow a similar expression pattern in response to changes in food intake (Akash et al., 2014). The increased expression of *cart2b* observed in high-density does not support the general anorexigenic role of *cart* genes, as less food availability in high-density is expected to reduce expression of anorexigenic genes. However, the tendency of *cart2a* towards increased expression in low-density follows the expected anorexigenic role, thus consistent with a hypothesis that suggests a potential functional divergence for *cart2a* and *cart2b* paralogs in the hypothalamus of Atlantic salmon. Interestingly, previous studies of Atlantic salmon also revealed a similar unexpected expression pattern for *cart2b* paralog, where its hypothalamic expression has been found significantly higher in fasted groups in response to both short- and long-term fasting (Kalananthan et al., 2021, 2023). In these studies, the authors suggested a potential orexigenic role for *cart2b* in the hypothalamus of Atlantic salmon, which is uncommon across teleost fish and remained to be functionally elucidated. Moreover, in our study, *cart2b* expression was strongly correlated with the orexigenic gene, *agrp1*, but only in high-density. Taken together, these results suggest that the neo-functionalized *cart2b* has obtained a role that appears to be opposite to the function previously described for CART, and which may be a mechanism that helps salmon adapt to changing environmental conditions. Furthermore, both *agrp1* and *cart2b* expression levels were negatively correlated with body mass, but only under high-density conditions. The negative correlation with *agrp1* is consistent with expectations, as reduced food availability in such conditions would likely lead to lower body mass and an increase in *agrp1* expression, a gene known to stimulate appetite. In contrast, the negative correlation with *cart2b* is unexpected for a gene that encodes an anorexigenic peptide. This finding aligns with previous observation that *cart2b* may have undergone functional divergence following genome duplication in Atlantic salmon (Kalananthan et al., 2021, 2023) resulting in a modified or divergent role from its original function.

### 4.3. Density affects the expression of m^6^A RNA methylation writer and reader genes

Given that m^6^A RNA methylation has been shown to be a mechanism sensitive to environmental changes (Frapin et al., 2020; Yang et al., 2022), such as variation in food availability and stress, we investigated the impact of population density on the expression of genes coding for writers, erasers, and readers of m^6^A RNA methylation.

We found a reduction in the hypothalamic expression of a writer, *mettl3*, and a reader, *ythdf1* (*ythdf1-2* paralog), in high-density. This suggests the potential role of these factors in mediating density response to stress and/or food availability in hypothalamus of Atlantic salmon. In recent years, the function of m^6^A RNA methylation factors in mediating stress signals has gained significant interest, with discoveries pointing to their versatile, tissue-specific, and context-dependent roles (Wilkinson et al., 2021). *mettl3* is a well-characterized and highly conserved m^6^A writer, the expression of which was reduced in high-density compared to low-density. Decreased *mettl3* expression under acute stress was earlier reported in the hypothalamus of mice (Engel et al., 2018). Strikingly, it seems that the stress mediated role of *mettl3* in the central nervous system is also highly conserved as a similar role was proposed in the *Drosophila* brain (Perlegos et al., 2022). The second density-affected m^6^A RNA methylation marker, *ythdf1-2*, encodes an m^6^A reader protein that facilitates translation initiation (X. Wang et al., 2015). Interestingly, findings in mammals have already revealed that *ythdf1* plays a role in various stress signals such as oxidative stress, hypoxia, heat and inflammatory stresses (He et al., 2022; Lu et al., 2019; Wilkinson et al., 2021). A recent study in rainbow trout also found several *ythdf1* paralogs to be responsive to heat and inflammatory stresses in various tissues (Yu et al., 2023). Although, to our knowledge, no study has indicated its role in stress signals of social competition, a novel discovery has shown that Ythdf1 is the main mediator of microbiota-dependent brain-gut crosstalk in mice (Huang et al., 2023). This raises a topic for future research, to understand whether *ythdf1-2* plays a role in translating gut signals triggered by food availability to the hypothalamus in salmon kept in different densities. The answer to this question is particularly interesting because it has been already demonstrated that microbiota-dependent brain-gut crosstalk is essential for regulation of both appetite and body temperature (Gabanyi et al., 2022).

### 4.4. Interconnection of stress, appetite regulation and m^6^A RNA methylation mechanism

In this study, several lines of evidence suggest that the processes/mechanisms targeted are interrelated. We observed significant correlations between *nr3c* paralogs and several m^6^A RNA methylation actors in both low- and high-density conditions. Notably, the strongest correlation was found between the gene coding for the main glucocorticoid receptor, *nr3c1,* and the m6A eraser *fto1,* and we also detected a negative correlation between *nr3c1* expression and the percentage of m^6^A RNA modifications only in the low-density condition. Interestingly, an association between *nr3c1* and *fto1,* has also been reported in recent studies, suggesting a dynamic and reciprocal regulation between FTO and NR3C1, mediated by m^6^A modifications (Roy et al., 2022; T. Wu et al., 2023). Moreover, our data suggest that this regulatory mechanism responds to the environment. Under high-density condition, while a correlation between *fto1* and *nr3c1* was still observed, there was no significant correlation between *nr3c1* and m^6^A methylation levels. This suggests that external factors, such as density-related stress, may alter the m^6^A-mediated regulation of these genes. This highlights the potential for a more complex, condition-specific interaction between *nr3c1* and *fto* that could influence m^6^A modifications and their downstream effects on gene expression.

Although we did not find any clear density-related difference in m^6^A RNA methylation levels, we cannot exclude the possibility that it is affected by density. Indeed, based on our results, this can be argued by (1) the effect of density on the expression of the gene encoding the core of the writing complex, *mettl3*, and (2) the effect of density on genes previously shown to be direct targets of m^6^A RNA methylation, such as CART mRNA in mice (Song et al., 2020; P.-F. Wu et al., 2021) and *crh* in chicken (Yang et al., 2022). Furthermore, we observed that *cart2b* was correlated with several m^6^A RNA readers under low-density conditions, including paralogs of genes involved in alternative splicing (*ythdc1.2*), in fate transition (*ythdc2.1*), in mRNA clearance (*ythdf2.2*) and translation (*ythdf3*.1) (Li et al., 2017; Liao et al., 2018). This suggests that *cart2b* may be regulated by a variety of m6A-modifying proteins, which play important roles in post-transcriptional processes. However, further investigation is required to understand what the mechanism behind the observed difference in *cart2b* expression between the two densities.

### 4.5. Gene expression associations with maturation locus genotypes

In Atlantic salmon, two loci near/across the genes *vgll3* and *six6* have been associated with sea age at maturity (Barson et al., 2015; Sinclair-Waters et al., 2020). Investigating whether genotypes at these loci influence gene expression in response to environmental factors such as population density, which can affect body mass, could provide insight into how genetic and environmental factors together regulate maturation.

In this study*, six6* genotype was associated with the expression of one gene related to stress, *crf1a1*, one gene related to appetite, *cart2a*, and three genes related to m^6^A RNA methylation, *mettl3, wtap* and *fto1*. For all these genes, the *six6*\*EL individuals had lower expression compared with *six6*\*EE and/or *six6*\*LL. Previous study of gene expression influenced by *six6* genotypes has only compared homozygous individuals (Ahi et al., 2024). In this study, we included heterozygotes and found notable differences in gene expression between heterozygous and homozygous individuals which may reflect a potential heterozygote advantage that warrants further investigation (Hedrick, 2012). As *six6* is coding for a transcription factor, it has the potential to directly regulate the expression of other genes. In addition, it remains an interesting area for future research to investigate whether six6 might influence gene expression through m^6^A methylation mechanisms. Although no differences in overall m^6^A RNA methylation levels were found between the *six6* genotypes, a more targeted analysis focusing on m^6^A methylation at the gene-specific level - particularly for genes affected by the *six6* genotype - would provide more informative insights.

We also found lower hypothalamic expression of *crf1b2* in the high-density compared to low-density condition, specifically in individuals with the *vgll3*\*LL genotype. It has been shown in Atlantic salmon that *crf1b2* paralogue is significantly induced in the telencephalon and hypothalamus in response to chasing stress (Lai et al., 2021). Moreover, in rainbow trout, CRF administration reduces aggressive behaviors but increases anxiety-like behaviors in a dose-dependent manner, suggesting a dual role of CRF1 in regulating both aggression and anxiety in the brain (Backström et al., 2011). Similarly, in rodents, lower doses of CRF can induce aggression, while higher doses tend to trigger anxiety-like responses (Hostetler & Ryabinin, 2013). Notably, a recent study showed that Atlantic salmon with the *vgll3*\*LL genotype exhibit higher aggression than those with the *vgll3*\*EE genotype (Bangura et al., 2022). This suggests that individuals with different *vgll3* genotypes may show distinct behavioral responses under stress, such as density pressure. It is possible that *vgll3* influences aggression in salmon by differentially regulating *crf1b2* in the hypothalamus, where the balance between aggression and anxiety-like behaviors may shift more drastically in *vgll3*\*LL individuals under high-density conditions due to altered *crf1b2* expression. Although no direct regulatory connection between *vgll3* (or the Hippo pathway) and *crf* genes has been identified in any species, the Hippo-vgll3 signal may indirectly modulate *CRF* gene expression through crosstalk with stress-related pathways, such as the glucocorticoid pathway, which acts upstream of *CRF*genes (Stepan et al., 2018).

## 5. Conclusions

Due to the importance of Atlantic salmon in aquaculture, the effects of density on gene expression have mainly been studied under farmed conditions (Conde-Sieira et al., 2010; Wunderink et al., 2011). Therefore, studies such as our that use near-natural densities can shed light on gene expression responses in more natural conditions. In this study, we found that the expression of genes related to stress, appetite and m6A RNA methylation were affected by density under semi-natural conditions. Our results suggest an interconnection between these three pathways. In addition, both *six6* and *vgll3* genotypes, which are associated with sea age at maturity in Atlantic salmon, appear to influence the expression of some of the genes studied. An intriguing finding regarding the *vgll3* genotype is that it may influence anxiety-related genes in a density-independent manner.

Our results have broader implications for other species, especially those with comparable ecological or evolutionary pressures, by highlighting how environmental factors and genetic variation interact to influence gene expression. Furthermore, the observed interconnection between the stress and appetite regulation, and m6A RNA methylation, which are well-conserved mechanisms across vertebrates (Ahi & Singh, 2024; Denver, 2009; Rønnestad et al., 2017), provided a framework for understanding these relationships in other species. Finally, our study highlights the importance of a deeper investigation of paralogs in salmonids, and other species with recent genome duplication, to better understand their specific functions, especially in adapting to environmental changes.

## Author Contributions

Experimental design (TA, JMP, OL, PH), conducting experiment (TA, JMP, OL, CRP), genotyping (TA, JMP), molecular experiment design (MF, EPA), molecular experiment (LQ, JT, MF), statistical analysis (MF, TA), funding (TA, CRP), writing - original draft (MF, EPA), writing - review and editing (MF, TA, EPA, JMP, CRP, PH, JT, OL).

## Acknowledgments

Funding was provided to CRP by Academy of Finland (grant numbers 307593, 302873, 327255 and 342851), the University of Helsinki, and the European Research Council under the European Articles Union’s Horizon 2020 and Horizon Europe research and innovation programs (grant nos. 742312 and 101054307). Views and opinions expressed are however those of the author(s) only and do not necessarily reflect those of the European Union or the European Research Council Executive Agency. Neither the European Union nor the granting authority can be held responsible for them. Funding was provided to TA by the Research Council of Finland (grant nos. 328860, 353388, and 325964). We would like to thank the Bodossaki Foundation for their financial support, which enabled Joana Troka to realize her MSc studies at the University of Helsinki. We thank Matyas Paszko, Atakan Yildiz, Paul Bangura, Ari Leinonen, Lison Gueguen, Ronan O’Sullivan, Nikolai Piavchenko, and Tommi Junnonaho for their help with fish rearing and experiment.

## Data Accessibility and Benefit-Sharing

All the data generated in this study are included in this article as supplementary file.

## Notes

### Competing Interest Statement

The authors have declared no competing interest.

### Summary of Updates

A typo in the author contributions part has been corrected in this new version, where in the molecular experiment TA has been corrected for JT.

