## Supplementary Figures 1-2 for "Density-dependent expression of epitranscriptomic, stress, and appetite regulating genes in Atlantic salmon"

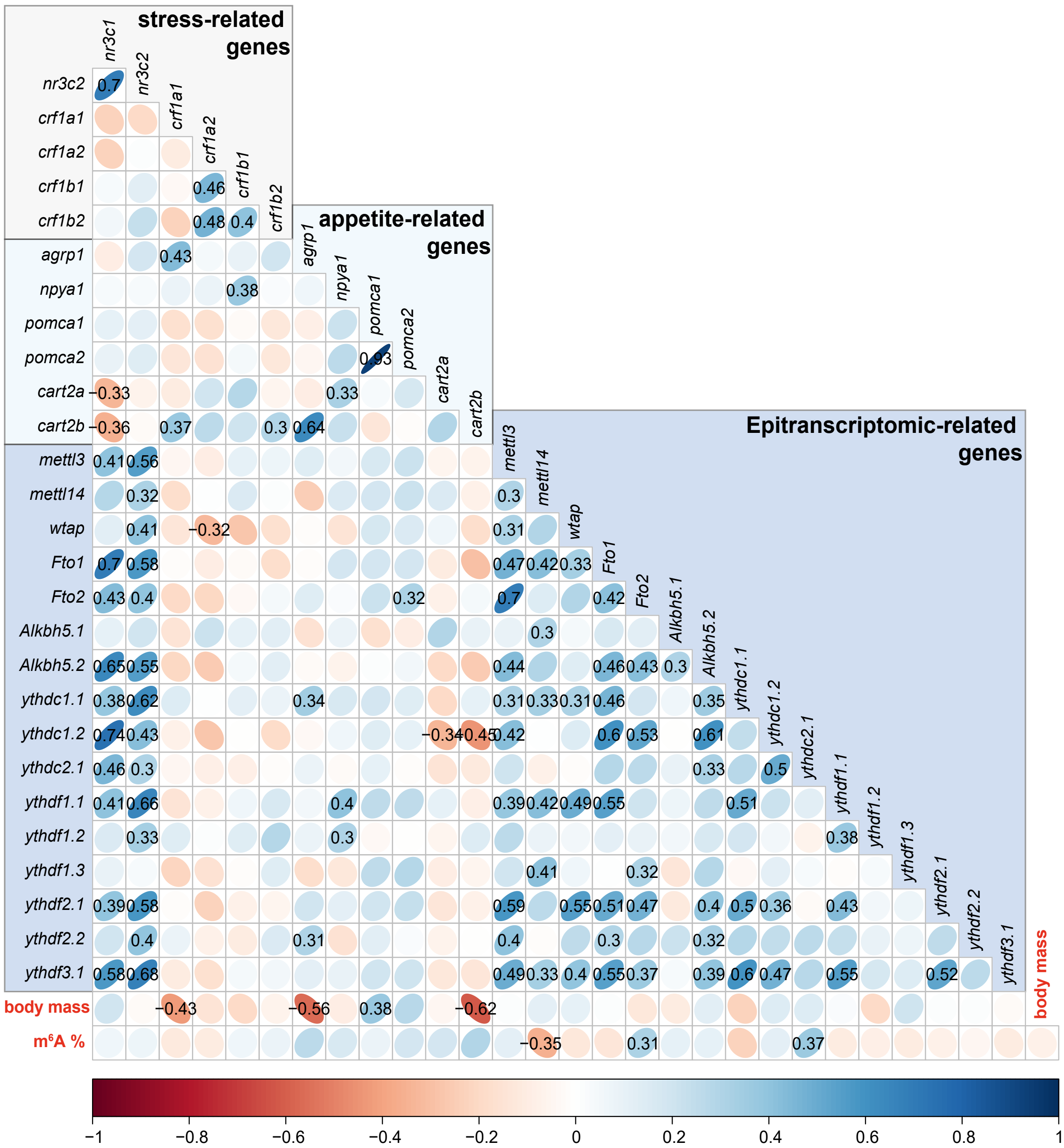


**Supplementary figure 1:** Pearson correlation results between the model residuals of gene expression, body mass and RNA m$^6$A methylation percentage in the high -density condition. The values shown in the graph correspond to the Pearson correlation coefficient of the significant correlation (p-value > 0.05).

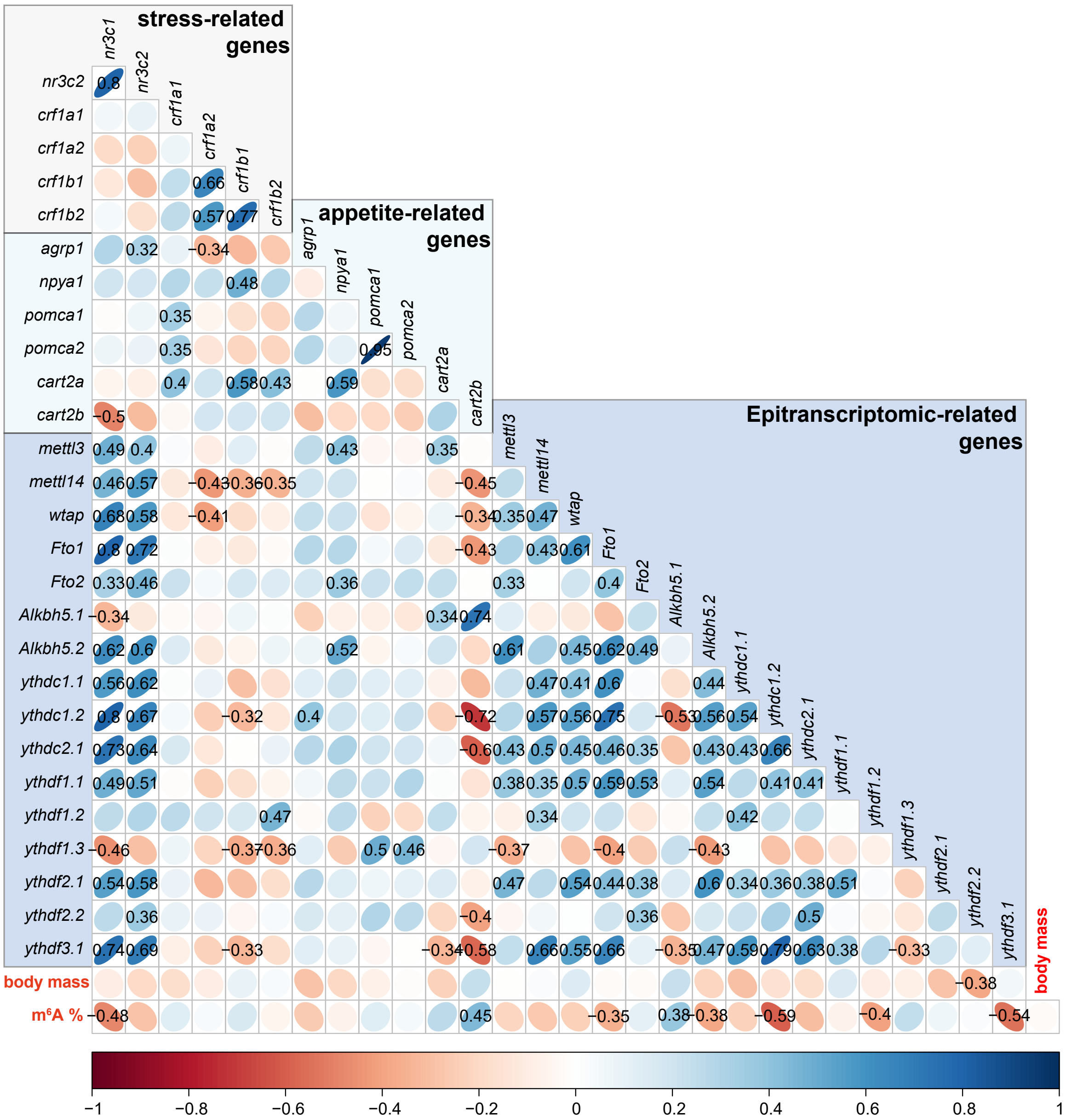


**Supplementary figure 2:** Pearson correlaotin results between the model residuals of gene expression, body mass and RNA m$^6$A methylaotin percentage in the low-density condition. The values shown in the graph correspond to the Pearson correlation coefficient of the significant correlation (p-value > 0.05).
