## Supplementary table 1-4 for "Density-dependent expression of epitranscriptomic, stress, and appetite regulating genes in Atlantic salmon"

**Supplementary table 1:** Number of fish and densities at important time points during rearing in semi-natural environmental conditions.

| | | **Density** | | | | | |
|---|---|---|---|---|---|---|---|
| | | **Low-density** | | | **High-density** | | |
| | | **Family**[1] | | | | | |
| **Date** | **Event** | **F3** | **F5** | **F8** | **F3** | **F5** | **F8** |
| 01 June 2021 | Fish transferred to stream channels (39 $m^2$) | 48 | 48 | 48 | 112 | 112 | 112 |
| | Density ($m^2$) | 1.21 | 1.21 | 1.21 | 2.84 | 2.84 | 2.84 |
| 24 June 2021 | Number of dead fish after peak outbreak | 27 | 9 | 15 | 86 | 33 | 52 |
| | Extra fish added (to enhance for density effects)[3] | | | | 74 | 22 | 41 |
| | Total fish | 21 | 39 | 33 | 100 | 111 | 101 |
| | Density ($m^2$) | 0.54 | 1.03 | 0.85 | 2.56 | 2.85 | 2.59 |
| 12 August 2021 | Transfer to the round streams (39.5 m2) | | | | | | |
| | Original fish | 15 | 16 | 21 | 18 | 51 | 47 |
| | Extra fish | | | | 73 | 20 | 35 |
| | Total fish | 15 | 16 | 21 | 91 | 71 | 82[2] |
| | Density ($m^2$) | 0.38 | 0.41 | 0.53 | 2.30 | 1.80 | 2.08 |
| 25 April 2022 | Final data count sand sampling | | | | | | |
| | Original fish | 13 | 17 | 20 | 17 | 47 | 42 |
| | Extra fish | | | | 64 | 17 | 33 |
| | Total fish (density / $m^2$) | 14 | 17 | 20 | 81 | 64 | 75 |
| | Density ($m^2$) | 0.35 | 0.43 | 0.51 | 2.05 | 1.62 | 1.90 |

1. Each family and density combination was reared in separate stream channels.
2. 12 fish was killed during transfer to the round streams.
3. Extra stream channel was supplemented by extra fish from the same family, which were reared in the same indoor tanks prior to the transfer of experimental fish to the natural stream channels.

**Supplementary table 2:** Number of fish used by family, sex and genotype, for the weight and the gene expression analysis.

| Locus | Sex | Genotypes | Density | | | | | |
|---|---|---|---|---|---|---|---|---|
| | | | Low-density | | | High-density | | |
| | | | Family | | | | | |
| | | | F3 | F5 | F8 | F3 | F5 | F8 |
| *vgll3* | Females | EE | 3 | 4 | 2 | 1 | 3 | 4 |
| | | EL | 1 | 0 | 3 | 1 | 1 | 3 |
| | | LL | 1 | 3 | 2 | 2 | 3 | 3 |
| | Males | EE | 0 | 1 | 2 | 0 | 4 | 4 |
| | | EL | 3 | 2 | 1 | 0 | 3 | 4 |
| | | LL | 3 | 2 | 4 | 2 | 5 | 1 |
| *six6* | Females | EE | 1 | 3 | 3 | 0 | 2 | 3 |
| | | EL | 1 | 2 | 1 | 2 | 2 | 2 |
| | | LL | 3 | 2 | 3 | 2 | 3 | 5 |
| | Males | EE | 2 | 2 | 3 | 1 | 4 | 3 |
| | | EL | 1 | 0 | 2 | 1 | 3 | 3 |
| | | LL | 3 | 3 | 2 | 0 | 5 | 3 |

**Supplementary table 3:** Full name, primer sequences and chromosomal location of the genes studied.

| | Full gene name | Ensembl ID | Forward primers (5' - 3') | Reverse primers (5' - 3') | Chr. |
|---|---|---|---|---|---|
| **Housekeeping genes** | | | | | |
| *ef1a* | elongation factor 1-alpha | ENSSSAG00000062937 | GCCTACCCTCCCCTTGGC | GTCACCTTGCCAGTGCTGG | ssa14 |
| *hprt1* | hypoxanthine phosphoribosyltransferase 1 | ENSSSAG00000039030 | GACTCATCCTTGACAGGACAGAGAG | CTTGAGCACGCAGAGAGCC | ssa09 |
| **Stress-related genes** | | | | | |
| *nr3c1* | Nuclear Receptor Subfamily 3 Group C Member 1 | ENSSSAG00000062169 | CCAGCAGCTTTGCCAGTTCA | GACAGATCTTATGGGCGGTCC | ssa05 |
| *nr3c2* | Nuclear Receptor Subfamily 3 Group C Member 2 | ENSSSAG00000087524 | GCCAGACAGCATGTCAAGCG | GCTGCCGCATGTAACAACC | ssa04 |
| *crf1a1* | Corticotropin-Releasing Factor | ENSSSAG00000079049 | GGAGCACTTGATCCATTCCACAATC | GATTTATTCGACAATGAGGACTGGGG | ssa03 |
| *crf1a2* | Corticotropin-Releasing Factor | ENSSSAG00000069223 | GGTCCATCCATCCCACGATCTA | AGGAGATGTGTTCGGCGATGA | ssa14 |
| *crf1b1* | Corticotropin-Releasing Factor | ENSSSAG00000052094 | CTCCACCGCTCCACAGCC | CCTCGGGGTGCATGACTTTC | ssa19 |
| *crf1b2* | Corticotropin-Releasing Factor | ENSSSAG00000080751 | CGCCACCGTTCCACATCAC | GGAGCCCTCTGGATACATGC | ssa29 |
| **Appetite-related genes** | | | | | |
| *agrp1* | agouti-related protein | ENSSSAG00000065164 | GGAATCCTACGATGAGGATGTTGCT | GACAGGACTGCTGGTGGG | ssa11 |
| *cart2a* | Cocaine- and amphetamine- | ENSSSAG00000015472 | AGTTCCCACTTGCGACGTG | AACACAATGATAGGGGATTGAATCCA | ssa10 |
| *cart2b* | regulated transcript | ENSSSAG00000047899 | CCCTACGTGCGACGTTGG | TTCAATTCACAAGCACTTGAGCAGA | ssa11 |
| *pomca1* | proopiomelanocortin | ENSSSAG00000004874 | TACTTTTGAAACAGCGTGACGATGC | TGCACTCCAGGATGCTGTTC | ssa09 |
| *pomca2* | proopiomelanocortin | ENSSSAG00000070765 | GAAGATTTGGCGACAGGCGAAG | CAGAGCTGAGGTCATGACAGC | ssa01 |
| *npya1* | neuropeptide Y | ENSSSAG00000040508 | GGTCAAACCCGAAACCCCC | CCTCTTCCCATACCTCTGCCT | ssa14 |
| **Epitranscriptomic-related genes** | | | | | |
| *mettl3* | methyltransferase 3 | ENSSSAG00000066322 | TCGTGCTCAGGTCCAGGAG | GATGATGCGTCGGAAGTGCAG | ssa18 |
| *mettl14* | methyltransferase 14 | ENSSSAG00020018770 | GAATAGTCGCCTACAGGAGATACGG | ACTCTCTGCTCCCAACTGTTGAG | NA |
| *wtap* | WT1 associated protein | ENSSSAG00000074636 | GGCTGAAGGAGTCTGAGGAGAAG | TGGATCTGGGTTGTACACTCTTGC | ssa15 |
| *alkbh5-1* | alkB homolog 5 | ENSSSAG00000058517 | ACAGTGATCCTGAGGCGAGG | CAACCACCTCATTTATCTTACCCTCG | ssa12 |
| *alkbh5-2* | alkB homolog 5 | ENSSSAG00000003588 | ATCCTGTGGAGTCCCGTGAAC | CAACCACCTCATTTATCTTCCCCTCA | ssa02 |
| *fto-1* | fat mass and obesity-associated gene | ENSSSAG00000053871 | ACCTGAGAGATGACCTGAACAGG | GAACACTCCGCCACTCTGTG | ssa26 |
| *fto-2* | fat mass and obesity-associated gene | ENSSSAG00000076575 | CCTACTACCTGAGAGGTACTAGAGCA | TCTATGGGTCAGTAGGGACACAGT | ssa11 |
| *ythdc1-1* | YTH N6-methyladenosine RNA binding protein C1 | ENSSSAG00000072116 | CCTATGAGAGATGGCGGGGT | ATCGGAGGACTCAGCTTCTCG | ssa13 |
| *ythdc1-2* | YTH N6-methyladenosine RNA binding protein C1 | ENSSSAG00000048322 | CTGGTGGTGGGCAGAGGG | CTGGGACGTATCAGCTTCTCTGTAG | ssa01 |
| *ythdc2* | YTH N6-methyladenosine RNA binding protein C2 | ENSSSAG00000058261 | ACAGCCACAGACAGGTCCTC | CCCTTGTCCCTGCTGCCA | ssa11 |
| *ythdf1-1* | YTH N6-methyladenosine RNA binding protein F1 | ENSSSAG00000048494 | GCAGGTGATGAAGATCATCGTGG | GCCAAGGATAAATGTCTTCTACTGCTG | ssa13 |

| | | | | | |
|---|---|---|---|---|---|
| *ythdf1-2* | YTH N6-methyladenosine RNA binding protein F1 | ENSSSAG00000055205 | GGTGCTCAAGATCATCGCCACT | GGACCCTCTCTGTATTGGAGCT | ssa13 |
| *ythdf1-3* | YTH N6-methyladenosine RNA binding protein F1 | ENSSSAG00000071507 | GTGCTGAAGATCATCGTGGGC | CTATTGTGGTTTACTTCGGTTTTGGTG | ssa15 |
| *ythdf2-1* | YTH N6-methyladenosine RNA binding protein F2 | ENSSSAG00000008511 | CAATGCCTATACGGCCATGTCG | GTAAGGCATGGGAGGATCTCCA | ssa01 |
| *ythdf2-2* | YTH N6-methyladenosine RNA binding protein F2 | ENSSSAG00000002258 | CAATGCGTATACGGCCATGTCA | TAAGGCATGGGGGGGTCG | ssa09 |
| *ythdf3* | YTH N6-methyladenosine RNA binding protein F3 | ENSSSAG00000056509 | TCAGGAGGTGCCTCTGGAG | TCTATTCCGCTCCCTCCGC | ssa29 |

**Supplementary table 4:** Summary of Analysis of Variance (ANOVA) results.

| | Genes | Terms | Sum Sq | Mean Sq | NumDF | DenDF | F value | Pr(>F) | Adjusted Pr(>F) |
|---|---|---|---|---|---|---|---|---|---|
| **Stress-related genes** | | **Density** | 0.133 | 0.133 | 1 | 79.0 | 1.095 | 0.299 | 0.492 |
| | | **Sex** | 0.266 | 0.266 | 1 | 77.8 | 2.182 | 0.144 | 0.402 |
| | | **six6** | 0.706 | 0.353 | 2 | 78.2 | 2.899 | 0.061 | 0.155 |
| | ***nr3c1*** | **vgll3** | 0.287 | 0.144 | 2 | 78.4 | 1.180 | 0.313 | 0.398 |
| | | **Density:Sex** | 0.158 | 0.158 | 1 | 78.3 | 1.299 | 0.258 | 0.481 |
| | | **Density:six6** | 0.116 | 0.058 | 2 | 77.8 | 0.477 | 0.622 | 0.792 |
| | | **Density:vgll3** | 0.060 | 0.030 | 2 | 78.1 | 0.246 | 0.783 | 0.894 |
| | | **Density** | 0.292 | 0.292 | 1 | 79.4 | 5.244 | 0.025 | 0.069 |
| | | **Sex** | 0.011 | 0.011 | 1 | 77.5 | 0.195 | 0.660 | 0.845 |
| | | **six6** | 0.060 | 0.030 | 2 | 78.1 | 0.539 | 0.585 | 0.755 |
| | ***nr3c2*** | **vgll3** | 0.238 | 0.119 | 2 | 78.6 | 2.136 | 0.125 | 0.269 |
| | | **Density:Sex** | 0.113 | 0.113 | 1 | 78.3 | 2.030 | 0.158 | 0.369 |
| | | **Density:six6** | 0.079 | 0.039 | 2 | 77.5 | 0.704 | 0.498 | 0.732 |
| | | **Density:vgll3** | 0.058 | 0.029 | 2 | 78.0 | 0.519 | 0.597 | 0.880 |
| | | **Density** | 3.737 | 3.737 | 1 | 79.3 | 8.598 | 0.004 | **0.025** |
| | | **Sex** | 0.043 | 0.043 | 1 | 77.8 | 0.098 | 0.755 | 0.845 |
| | | **six6** | 5.889 | 2.945 | 2 | 78.2 | 6.775 | 0.002 | **0.018** |
| | ***crf1a1*** | **vgll3** | 1.309 | 0.655 | 2 | 78.6 | 1.506 | 0.228 | 0.336 |
| | | **Density:Sex** | 0.913 | 0.913 | 1 | 78.3 | 2.101 | 0.151 | 0.369 |
| | | **Density:six6** | 1.461 | 0.731 | 2 | 77.8 | 1.681 | 0.193 | 0.684 |
| | | **Density:vgll3** | 0.857 | 0.429 | 2 | 78.2 | 0.986 | 0.378 | 0.705 |
| | | **Density** | 0.153 | 0.153 | 1 | 81.0 | 0.228 | 0.635 | 0.720 |
| | | **Sex** | 3.696 | 3.696 | 1 | 81.0 | 5.490 | 0.022 | 0.107 |
| | | **six6** | 0.333 | 0.166 | 2 | 81.0 | 0.247 | 0.782 | 0.878 |
| | ***crf1a2*** | **vgll3** | 2.392 | 1.196 | 2 | 81.0 | 1.777 | 0.176 | 0.308 |
| | | **Density:Sex** | 1.909 | 1.909 | 1 | 81.0 | 2.836 | 0.096 | 0.299 |
| | | **Density:six6** | 1.103 | 0.552 | 2 | 81.0 | 0.820 | 0.444 | 0.732 |
| | | **Density:vgll3** | 2.240 | 1.120 | 2 | 81.0 | 1.664 | 0.196 | 0.705 |
| | | **Density** | 6.967 | 6.967 | 1 | 81.0 | 13.801 | 0.000 | **0.003** |
| | | **Sex** | 3.137 | 3.137 | 1 | 81.0 | 6.215 | 0.015 | 0.107 |
| | | **six6** | 2.638 | 1.319 | 2 | 81.0 | 2.613 | 0.079 | 0.185 |
| | ***crf1b1*** | **vgll3** | 4.014 | 2.007 | 2 | 81.0 | 3.976 | 0.023 | 0.090 |
| | | **Density:Sex** | 2.310 | 2.310 | 1 | 81.0 | 4.576 | 0.035 | 0.231 |
| | | **Density:six6** | 1.742 | 0.871 | 2 | 81.0 | 1.726 | 0.184 | 0.684 |
| | | **Density:vgll3** | 2.820 | 1.410 | 2 | 81.0 | 2.793 | 0.067 | 0.705 |
| | | **Density** | 0.461 | 0.461 | 1 | 80.5 | 4.314 | 0.041 | 0.104 |
| | | **Sex** | 1.029 | 1.029 | 1 | 78.6 | 9.622 | 0.003 | 0.075 |
| | ***crf1b2*** | **six6** | 0.867 | 0.433 | 2 | 79.2 | 4.053 | 0.021 | 0.087 |
| | | **vgll3** | 1.081 | 0.540 | 2 | 80.0 | 5.051 | 0.009 | 0.074 |
| | | **Density:Sex** | 0.484 | 0.484 | 1 | 79.6 | 4.529 | 0.036 | 0.231 |

| | | | | | | | | | |
|---|---|---|---|---|---|---|---|---|---|
| | | **Density:six6** | 0.195 | 0.098 | 2 | 78.5 | 0.912 | 0.406 | 0.732 |
| | | **Density:vgll3** | 1.529 | 0.764 | 2 | 79.2 | 7.147 | 0.001 | **0.039** |
| **Appetite-related genes** | ***agrp1*** | **Density** | 26.474 | 26.474 | 1 | 78.8 | 51.640 | 0.000 | **0.000** |
| | | **Sex** | 0.005 | 0.005 | 1 | 78.0 | 0.011 | 0.918 | 0.918 |
| | | **six6** | 0.955 | 0.478 | 2 | 78.3 | 0.932 | 0.398 | 0.619 |
| | | **vgll3** | 0.718 | 0.359 | 2 | 78.4 | 0.701 | 0.499 | 0.583 |
| | | **Density:Sex** | 0.039 | 0.039 | 1 | 78.3 | 0.076 | 0.783 | 0.847 |
| | | **Density:six6** | 2.503 | 1.251 | 2 | 78.0 | 2.441 | 0.094 | 0.684 |
| | | **Density:vgll3** | 0.192 | 0.096 | 2 | 78.2 | 0.187 | 0.830 | 0.894 |
| | ***cart2a*** | **Density** | 2.007 | 2.007 | 1 | 81.0 | 6.511 | 0.013 | **0.050** |
| | | **Sex** | 1.347 | 1.347 | 1 | 81.0 | 4.369 | 0.040 | 0.159 |
| | | **six6** | 5.783 | 2.891 | 2 | 81.0 | 9.380 | 0.000 | **0.006** |
| | | **vgll3** | 0.325 | 0.162 | 2 | 81.0 | 0.527 | 0.593 | 0.638 |
| | | **Density:Sex** | 0.675 | 0.675 | 1 | 81.0 | 2.188 | 0.143 | 0.369 |
| | | **Density:six6** | 0.023 | 0.011 | 2 | 81.0 | 0.037 | 0.964 | 0.964 |
| | | **Density:vgll3** | 0.419 | 0.209 | 2 | 81.0 | 0.679 | 0.510 | 0.793 |
| | ***cart2b*** | **Density** | 3.122 | 3.122 | 1 | 80.5 | 12.666 | 0.001 | **0.004** |
| | | **Sex** | 0.219 | 0.219 | 1 | 78.7 | 0.888 | 0.349 | 0.678 |
| | | **six6** | 0.497 | 0.249 | 2 | 79.3 | 1.009 | 0.369 | 0.619 |
| | | **vgll3** | 0.823 | 0.411 | 2 | 80.1 | 1.669 | 0.195 | 0.321 |
| | | **Density:Sex** | 0.701 | 0.701 | 1 | 79.7 | 2.842 | 0.096 | 0.299 |
| | | **Density:six6** | 0.339 | 0.169 | 2 | 78.6 | 0.687 | 0.506 | 0.732 |
| | | **Density:vgll3** | 0.114 | 0.057 | 2 | 79.3 | 0.232 | 0.794 | 0.894 |
| | ***pomca1*** | **Density** | 0.343 | 0.343 | 1 | 81.0 | 0.243 | 0.623 | 0.720 |
| | | **Sex** | 1.182 | 1.182 | 1 | 81.0 | 0.838 | 0.363 | 0.678 |
| | | **six6** | 3.567 | 1.783 | 2 | 81.0 | 1.264 | 0.288 | 0.538 |
| | | **vgll3** | 15.083 | 7.542 | 2 | 81.0 | 5.345 | 0.007 | 0.074 |
| | | **Density:Sex** | 4.752 | 4.752 | 1 | 81.0 | 3.368 | 0.070 | 0.299 |
| | | **Density:six6** | 4.964 | 2.482 | 2 | 81.0 | 1.759 | 0.179 | 0.684 |
| | | **Density:vgll3** | 3.240 | 1.620 | 2 | 81.0 | 1.148 | 0.322 | 0.705 |
| | ***pomca2*** | **Density** | 1.046 | 1.046 | 1 | 81.0 | 0.651 | 0.422 | 0.591 |
| | | **Sex** | 0.845 | 0.845 | 1 | 81.0 | 0.525 | 0.471 | 0.759 |
| | | **six6** | 2.129 | 1.064 | 2 | 81.0 | 0.662 | 0.518 | 0.726 |
| | | **vgll3** | 17.278 | 8.639 | 2 | 81.0 | 5.375 | 0.006 | 0.074 |
| | | **Density:Sex** | 2.931 | 2.931 | 1 | 81.0 | 1.824 | 0.181 | 0.389 |
| | | **Density:six6** | 6.925 | 3.463 | 2 | 81.0 | 2.154 | 0.123 | 0.684 |
| | | **Density:vgll3** | 3.673 | 1.836 | 2 | 81.0 | 1.143 | 0.324 | 0.705 |
| | ***npya1*** | **Density** | 5.097 | 5.097 | 1 | 81.0 | 5.868 | 0.018 | 0.055 |
| | | **Sex** | 6.881 | 6.881 | 1 | 81.0 | 7.921 | 0.006 | 0.086 |
| | | **six6** | 5.764 | 2.882 | 2 | 81.0 | 3.318 | 0.041 | 0.128 |
| | | **vgll3** | 8.378 | 4.189 | 2 | 81.0 | 4.822 | 0.011 | 0.074 |
| | | **Density:Sex** | 7.126 | 7.126 | 1 | 81.0 | 8.203 | 0.005 | 0.089 |
| | | **Density:six6** | 0.462 | 0.231 | 2 | 81.0 | 0.266 | 0.767 | 0.835 |

| | | | | | | | | | |
|---|---|---|---|---|---|---|---|---|---|
| | | **Density:vgll3** | 3.830 | 1.915 | 2 | 81.0 | 2.205 | 0.117 | 0.705 |
| **Epitranscriptomic-related genes** | *mettl3* | **Density** | 0.439 | 0.439 | 1 | 81.0 | 7.554 | 0.007 | **0.034** |
| | | **Sex** | 0.024 | 0.024 | 1 | 81.0 | 0.405 | 0.526 | 0.759 |
| | | **six6** | 0.989 | 0.495 | 2 | 81.0 | 8.520 | 0.000 | **0.006** |
| | | **vgll3** | 0.335 | 0.167 | 2 | 81.0 | 2.884 | 0.062 | 0.178 |
| | | **Density:Sex** | 0.012 | 0.012 | 1 | 81.0 | 0.205 | 0.652 | 0.827 |
| | | **Density:six6** | 0.149 | 0.074 | 2 | 81.0 | 1.281 | 0.283 | 0.732 |
| | | **Density:vgll3** | 0.102 | 0.051 | 2 | 81.0 | 0.877 | 0.420 | 0.735 |
| | *mettl14* | **Density** | 0.351 | 0.351 | 1 | 78.9 | 2.919 | 0.091 | 0.197 |
| | | **Sex** | 0.683 | 0.683 | 1 | 78.0 | 5.675 | 0.020 | 0.107 |
| | | **six6** | 0.932 | 0.466 | 2 | 78.3 | 3.874 | 0.025 | 0.087 |
| | | **vgll3** | 0.322 | 0.161 | 2 | 78.5 | 1.337 | 0.269 | 0.376 |
| | | **Density:Sex** | 0.008 | 0.008 | 1 | 78.3 | 0.065 | 0.800 | 0.847 |
| | | **Density:six6** | 0.646 | 0.323 | 2 | 78.0 | 2.685 | 0.075 | 0.684 |
| | | **Density:vgll3** | 0.261 | 0.131 | 2 | 78.3 | 1.086 | 0.343 | 0.705 |
| | *wtap* | **Density** | 0.121 | 0.121 | 1 | 81.0 | 1.018 | 0.316 | 0.492 |
| | | **Sex** | 0.168 | 0.168 | 1 | 81.0 | 1.417 | 0.237 | 0.604 |
| | | **six6** | 1.379 | 0.689 | 2 | 81.0 | 5.808 | 0.004 | **0.031** |
| | | **vgll3** | 0.012 | 0.006 | 2 | 81.0 | 0.049 | 0.953 | 0.988 |
| | | **Density:Sex** | 0.073 | 0.073 | 1 | 81.0 | 0.617 | 0.434 | 0.667 |
| | | **Density:six6** | 0.072 | 0.036 | 2 | 81.0 | 0.303 | 0.740 | 0.835 |
| | | **Density:vgll3** | 0.002 | 0.001 | 2 | 81.0 | 0.007 | 0.993 | 0.993 |
| | *alkbh5-1* | **Density** | 0.584 | 0.584 | 1 | 81.0 | 1.678 | 0.199 | 0.348 |
| | | **Sex** | 0.009 | 0.009 | 1 | 81.0 | 0.027 | 0.870 | 0.918 |
| | | **six6** | 0.651 | 0.326 | 2 | 81.0 | 0.936 | 0.396 | 0.619 |
| | | **vgll3** | 1.098 | 0.549 | 2 | 81.0 | 1.578 | 0.213 | 0.331 |
| | | **Density:Sex** | 0.310 | 0.310 | 1 | 81.0 | 0.891 | 0.348 | 0.573 |
| | | **Density:six6** | 0.178 | 0.089 | 2 | 81.0 | 0.256 | 0.775 | 0.835 |
| | | **Density:vgll3** | 0.243 | 0.122 | 2 | 81.0 | 0.349 | 0.706 | 0.894 |
| | *alkbh5-2* | **Density** | 0.001 | 0.001 | 1 | 80.4 | 0.019 | 0.889 | 0.889 |
| | | **Sex** | 0.037 | 0.037 | 1 | 78.7 | 0.837 | 0.363 | 0.678 |
| | | **six6** | 0.351 | 0.176 | 2 | 79.3 | 4.022 | 0.022 | 0.087 |
| | | **vgll3** | 0.351 | 0.176 | 2 | 79.8 | 4.021 | 0.022 | 0.090 |
| | | **Density:Sex** | 0.002 | 0.002 | 1 | 79.5 | 0.054 | 0.817 | 0.847 |
| | | **Density:six6** | 0.102 | 0.051 | 2 | 78.6 | 1.172 | 0.315 | 0.732 |
| | | **Density:vgll3** | 0.132 | 0.066 | 2 | 79.2 | 1.508 | 0.228 | 0.705 |
| | *fto-1* | **Density** | 0.008 | 0.008 | 1 | 78.5 | 0.113 | 0.738 | 0.765 |
| | | **Sex** | 0.036 | 0.036 | 1 | 77.9 | 0.499 | 0.482 | 0.759 |
| | | **six6** | 0.759 | 0.379 | 2 | 78.1 | 5.247 | 0.007 | **0.041** |
| | | **vgll3** | 0.152 | 0.076 | 2 | 78.2 | 1.054 | 0.353 | 0.430 |
| | | **Density:Sex** | 0.097 | 0.097 | 1 | 78.1 | 1.340 | 0.251 | 0.481 |
| | | **Density:six6** | 0.316 | 0.158 | 2 | 77.9 | 2.184 | 0.119 | 0.684 |
| | | **Density:vgll3** | 0.143 | 0.072 | 2 | 78.1 | 0.991 | 0.376 | 0.705 |

| | | | | | | | | |
|---|---|---|---|---|---|---|---|---|
| *fto-2* | **Density** | 0.030 | 0.030 | 1 | 81.0 | 0.217 | 0.643 | 0.720 |
| | **Sex** | 0.469 | 0.469 | 1 | 81.0 | 3.404 | 0.069 | 0.214 |
| | **six6** | 0.062 | 0.031 | 2 | 81.0 | 0.226 | 0.798 | 0.878 |
| | **vgll3** | 0.564 | 0.282 | 2 | 81.0 | 2.046 | 0.136 | 0.272 |
| | **Density:Sex** | 0.019 | 0.019 | 1 | 81.0 | 0.138 | 0.711 | 0.829 |
| | **Density:six6** | 0.205 | 0.102 | 2 | 81.0 | 0.744 | 0.479 | 0.732 |
| | **Density:vgll3** | 0.414 | 0.207 | 2 | 81.0 | 1.502 | 0.229 | 0.705 |
| *ythdc1-1* | **Density** | 0.012 | 0.012 | 1 | 80.2 | 0.121 | 0.729 | 0.765 |
| | **Sex** | 0.010 | 0.010 | 1 | 77.3 | 0.098 | 0.755 | 0.845 |
| | **six6** | 0.463 | 0.231 | 2 | 78.2 | 2.276 | 0.109 | 0.236 |
| | **vgll3** | 0.581 | 0.290 | 2 | 79.3 | 2.857 | 0.063 | 0.178 |
| | **Density:Sex** | 0.036 | 0.036 | 1 | 78.8 | 0.358 | 0.551 | 0.772 |
| | **Density:six6** | 0.070 | 0.035 | 2 | 77.1 | 0.346 | 0.709 | 0.835 |
| | **Density:vgll3** | 0.057 | 0.028 | 2 | 78.1 | 0.280 | 0.757 | 0.894 |
| *ythdc1-2* | **Density** | 0.031 | 0.031 | 1 | 80.6 | 0.225 | 0.637 | 0.720 |
| | **Sex** | 0.168 | 0.168 | 1 | 78.4 | 1.227 | 0.271 | 0.633 |
| | **six6** | 0.217 | 0.109 | 2 | 79.1 | 0.795 | 0.455 | 0.671 |
| | **vgll3** | 0.724 | 0.362 | 2 | 80.3 | 2.649 | 0.077 | 0.196 |
| | **Density:Sex** | 0.427 | 0.427 | 1 | 79.8 | 3.123 | 0.081 | 0.299 |
| | **Density:six6** | 0.219 | 0.110 | 2 | 78.0 | 0.802 | 0.452 | 0.732 |
| | **Density:vgll3** | 0.189 | 0.095 | 2 | 79.1 | 0.692 | 0.503 | 0.793 |
| *ythdc2-1* | **Density** | 0.249 | 0.249 | 1 | 79.2 | 1.696 | 0.197 | 0.348 |
| | **Sex** | 0.064 | 0.064 | 1 | 77.6 | 0.438 | 0.510 | 0.759 |
| | **six6** | 0.871 | 0.436 | 2 | 78.1 | 2.965 | 0.057 | 0.155 |
| | **vgll3** | 0.193 | 0.096 | 2 | 78.4 | 0.656 | 0.522 | 0.584 |
| | **Density:Sex** | 0.633 | 0.633 | 1 | 78.2 | 4.309 | 0.041 | 0.231 |
| | **Density:six6** | 0.453 | 0.227 | 2 | 77.6 | 1.542 | 0.220 | 0.686 |
| | **Density:vgll3** | 0.043 | 0.021 | 2 | 78.0 | 0.146 | 0.865 | 0.897 |
| *ythdf1-1* | **Density** | 0.257 | 0.257 | 1 | 78.6 | 5.950 | 0.017 | 0.055 |
| | **Sex** | 0.007 | 0.007 | 1 | 77.8 | 0.154 | 0.696 | 0.845 |
| | **six6** | 0.018 | 0.009 | 2 | 78.1 | 0.204 | 0.816 | 0.878 |
| | **vgll3** | 0.388 | 0.194 | 2 | 78.2 | 4.505 | 0.014 | 0.079 |
| | **Density:Sex** | 0.002 | 0.002 | 1 | 78.1 | 0.037 | 0.848 | 0.848 |
| | **Density:six6** | 0.144 | 0.072 | 2 | 77.8 | 1.668 | 0.195 | 0.684 |
| | **Density:vgll3** | 0.101 | 0.050 | 2 | 78.0 | 1.170 | 0.316 | 0.705 |
| *ythdf1-2* | **Density** | 0.828 | 0.828 | 1 | 79.5 | 14.820 | 0.000 | **0.003** |
| | **Sex** | 0.301 | 0.301 | 1 | 77.5 | 5.387 | 0.023 | 0.107 |
| | **six6** | 0.007 | 0.003 | 2 | 78.1 | 0.061 | 0.941 | 0.941 |
| | **vgll3** | 0.263 | 0.132 | 2 | 78.6 | 2.358 | 0.101 | 0.236 |
| | **Density:Sex** | 0.032 | 0.032 | 1 | 78.3 | 0.570 | 0.452 | 0.667 |
| | **Density:six6** | 0.057 | 0.028 | 2 | 77.4 | 0.507 | 0.605 | 0.792 |
| | **Density:vgll3** | 0.227 | 0.113 | 2 | 78.0 | 2.032 | 0.138 | 0.705 |
| *ythdf1-3* | **Density** | 0.094 | 0.094 | 1 | 81.0 | 0.694 | 0.407 | 0.591 |

| | | | | | | | | |
|---|---|---|---|---|---|---|---|---|
| | **Sex** | 0.051 | 0.051 | 1 | 81.0 | 0.375 | 0.542 | 0.759 |
| | **six6** | 0.341 | 0.171 | 2 | 81.0 | 1.266 | 0.287 | 0.538 |
| | **vgll3** | 0.001 | 0.001 | 2 | 81.0 | 0.005 | 0.995 | 0.995 |
| | **Density:Sex** | 0.032 | 0.032 | 1 | 81.0 | 0.235 | 0.629 | 0.827 |
| | **Density:six6** | 0.257 | 0.128 | 2 | 81.0 | 0.953 | 0.390 | 0.732 |
| | **Density:vgll3** | 0.066 | 0.033 | 2 | 81.0 | 0.243 | 0.785 | 0.894 |
| ***ythdf2-1*** | **Density** | 0.156 | 0.156 | 1 | 81.0 | 1.852 | 0.177 | 0.348 |
| | **Sex** | 0.002 | 0.002 | 1 | 81.0 | 0.020 | 0.888 | 0.918 |
| | **six6** | 0.081 | 0.040 | 2 | 81.0 | 0.481 | 0.620 | 0.755 |
| | **vgll3** | 0.306 | 0.153 | 2 | 81.0 | 1.815 | 0.169 | 0.308 |
| | **Density:Sex** | 0.087 | 0.087 | 1 | 81.0 | 1.032 | 0.313 | 0.547 |
| | **Density:six6** | 0.110 | 0.055 | 2 | 81.0 | 0.653 | 0.523 | 0.732 |
| | **Density:vgll3** | 0.246 | 0.123 | 2 | 81.0 | 1.461 | 0.238 | 0.705 |
| ***ythdf2-2*** | **Density** | 0.064 | 0.064 | 1 | 81.0 | 0.255 | 0.615 | 0.720 |
| | **Sex** | 0.042 | 0.042 | 1 | 81.0 | 0.168 | 0.683 | 0.845 |
| | **six6** | 0.255 | 0.128 | 2 | 81.0 | 0.510 | 0.603 | 0.755 |
| | **vgll3** | 0.612 | 0.306 | 2 | 81.0 | 1.222 | 0.300 | 0.398 |
| | **Density:Sex** | 0.043 | 0.043 | 1 | 81.0 | 0.172 | 0.679 | 0.827 |
| | **Density:six6** | 0.044 | 0.022 | 2 | 81.0 | 0.088 | 0.916 | 0.950 |
| | **Density:vgll3** | 0.102 | 0.051 | 2 | 81.0 | 0.204 | 0.816 | 0.894 |
| ***ythdf3-1*** | **Density** | 0.311 | 0.311 | 1 | 81.0 | 3.926 | 0.051 | 0.119 |
| | **Sex** | 0.274 | 0.274 | 1 | 81.0 | 3.456 | 0.067 | 0.214 |
| | **six6** | 0.021 | 0.011 | 2 | 81.0 | 0.133 | 0.876 | 0.908 |
| | **vgll3** | 0.500 | 0.250 | 2 | 81.0 | 3.150 | 0.048 | 0.169 |
| | **Density:Sex** | 0.622 | 0.622 | 1 | 81.0 | 7.845 | 0.006 | 0.089 |
| | **Density:six6** | 0.125 | 0.062 | 2 | 81.0 | 0.787 | 0.459 | 0.732 |
| | **Density:vgll3** | 0.194 | 0.097 | 2 | 81.0 | 1.220 | 0.301 | 0.705 |
